# Impact of Sex and *APOE* Status on Spatial Navigation in Pre-symptomatic Alzheimer’s disease

**DOI:** 10.1101/287722

**Authors:** G. Coughlan, A. Coutrot, M. Khondoker, A. Minihane, H. Spiers, M. Hornberger

## Abstract

**INTRODUCTION:** Spatial navigation is emerging as a critical factor in identifying pre-symptomatic Alzheimer pathophysiology, with the impact of sex and *APOE* status on spatial navigation yet to be established.

**METHODS:** We estimate the effects of sex on navigation performance in 27,308 individuals (50-70 years [benchmark population]) by employing a novel game-based approach to cognitive assessment using *Sea Hero Quest*. The effects of *APOE* genotype and sex on game performance was further examined in a smaller lab-based cohort (*n* = 44).

**RESULTS:** Benchmark data showed an effect of sex on wayfinding distance, duration and path integration. Importantly in the lab cohort, performance on allocentric wayfinding levels was reduced in *ε4* carriers compared to *ε3* carriers, and effect of sex became negligible when *APOE* status was controlled for. To demonstrate the robustness of this effect and to ensure the quality of data obtained through unmonitored at-home use of the Sea Hero Quest game, post-hoc analysis was carried out to compare performance by the benchmark population to the monitored lab-cohort.

**DISCUSSION:** *APOE ε4* midlife carriers exhibit changes in navigation pattern before any symptom onset. This supports the move towards spatial navigation as an early cognitive marker and demonstrates for the first time how the utility of large-scale digital cognitive assessment may hold future promise for the early detection of Alzheimer’s disease. Finally, benchmark findings suggest that gender differences may need to be considered when determining the classification criteria for spatial navigational deficits in midlife adults.

## 1. Background

Spatial navigation is emerging as a highly promising cognitive biomarker for underlying Alzheimer’s disease pathophysiology ([1–4]). Indeed, the brain areas earliest affected by AD are key nodes in the spatial navigation network [5–7]. Increasing evidence suggest that such spatial navigation changes can be present before episodic memory deficits, which are the current gold standard to diagnosis of AD [8–10]. Thus, spatial navigation emerges as a potential cost effective cognitive biomarker that can detect AD in the pre-symptomatic (i.e. before memory deficits) stages ([11,12], which has significant implications for future diagnostics and treatment approaches.

However, little is still known about variation of spatial navigation processes on a population basis. For example, contemporary legend has attempted to explain why men and women navigate differently, although there is no existing population level navigation data to support such myths. Understanding this variation on a population level is important because sex has been known to impact on the incidence of AD, with the age-specific risk of AD being almost two-fold greater in women than men (17.2% versus 9.1% at age 65 years for example)[13,14]. Similarly, the impact of genetic-risk on spatial navigation is still being established. Emerging evidence suggests that *APOE ε4* carriers (heterozygous or homozygous) show altered spatial navigation compared to *APOE ε3* homozygous carriers [11,12]. But, it is not clear how sex might interact with *APOE* status to influence spatial navigation. Clearly, given the prevalence of the *APOE ε4* genotype (20-25% Caucasians), delineating the individual and interactive impact of sex and *APOE* status on is of high importance if spatial navigation will emerge as a new cognitive biomarker.

In the current study, we examine the effect of sex and APOE status on spatial navigation performance in midlife adults. Using the Sea Hero Quest spatial navigation APP, we examined the effect of sex on performance in the benchmark population cohort (*n*=27,308) and further examined the individual and joint effect of gender and APOE status using a lab cohort (*n*=44) that has been *APOE* genotyped. We hypothesise that men and women will differ on spatial navigation performance and this sex effect will interact with *APOE* status.

## 2. Methods

The population-based data was extracted the global Sea Hero Quest (SHQ) database (see below) that includes spatial navigation data on over 3 million people worldwide. The lab-based cohort was tested on the same SHQ App as the benchmark cohort; in addition, participants were *APOE* genotyped.

### 2.1 Population cohort

#### Data collection

The SHQ database was generated using a mobile-app based cognitive task that measures spatial navigation ability [27]. This app was developed in 2015 by our team and funded by Deutsche Telekom and Alzheimer’s Research UK. It was made available for free on the App Store and Play Store from May 2016 and since then over 3 million people have downloaded the App worldwide (www.seaheroquest.com). Participants were given the option to opt-in or –out of the data collection. If a participants’ response was to opt-in, their SHQ data was anonymised and stored securely by the T-Systems’ datacentre under the regulation of German data security law. The study was ethically approved by an Ethics Research Committee CPB/2013/015.

#### Game description

The game performance can be divided into two main domains: wayfinding and path integration (PI). In wayfinding levels (here 1, 2, 6, 8 and 11), players are initially presented with a map indicating start location and the location of several checkpoints to find in a set order. The two variables of interest are trajectory distance and duration taken to complete each level. In path integration levels (here 9 and 14), participants navigate along a river with bends to find a flare gun and then choose from a choice of three possible directions which is the correct direction back to the starting point along the Euclidean (Figure 1. i). Depending on their accuracy, players receive either one, two or three stars. Given that video gaming proficiency could bias performance by giving players familiar with similar games an advantage, spatial navigation scores on wayfinding levels within the database were normalised against the sum of participant scores on Level 1 and 2 which were classified as practice levels given that neither level requires navigation abilities.

**Figure 1.**
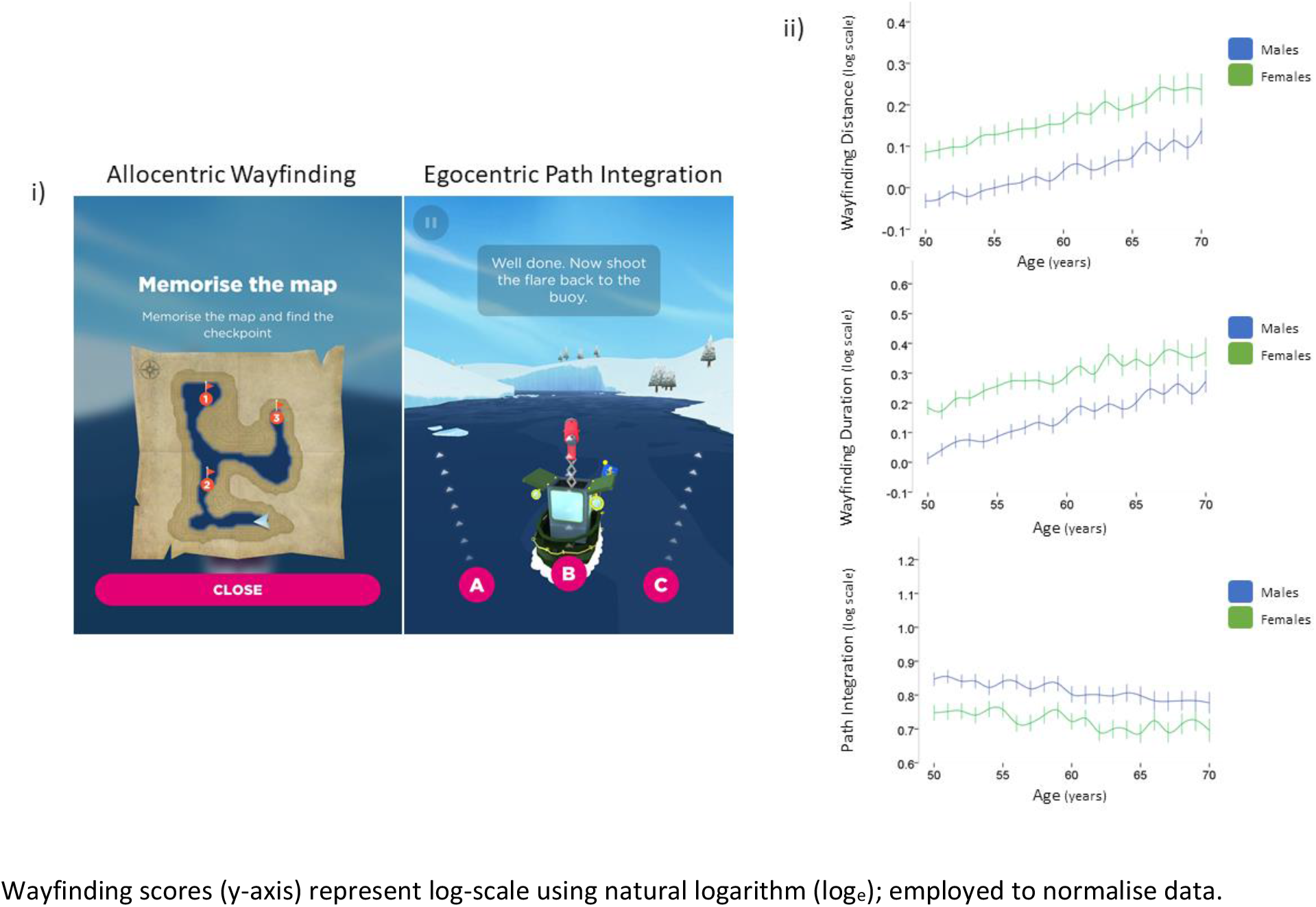
i) Example of map participants are invited to study before the start the wayfinding task (left). Example of starting direction choice presented to participants following path integration (right). ii) Sex and age on wayfinding distance, wayfinding duration and path integration performance for the benchmark population.

#### Study Population

Demographic information, specifically age, sex and nationality is available from the global SHQ database and were used for this research. We restricted the age range to 50-70 years as they are the most vulnerable age-group to develop AD in the next decade, and therefore the detection of spatial deficits may facilitate early detection of underlying AD pathology. We also restricted the data to the UK population only, to allow cross-comparison to our lab cohort. To provide a reliable estimate of spatial navigation ability, only participants who completed all above-mentioned levels were included, allowing us to examine wayfinding and path integration abilities in 13,461 men and 13,847 women from the UK (total *n* = 27,308).

### 2.2 Lab-based cohort

#### Data collection

A total of 120 participants between 50 and 75 years old were recruited to participate in the lab cohort study. Participants were screened for a history of psychiatric disorders, history of drug dependency disorders, significant relevant comorbidity or visual loss likely to interfere with the research protocol. If a participants’ ACE-III scores were lower than 85, they were classified as suspected mildly cognitively impaired and excluded from the study. Three participants who showed signs of frustration or emotional unease were also excluded. Ethical approval was obtained from the Faculty of Medicine and Health Sciences Ethics Committee at the University of East Anglia Reference 2016/2017 – 11.

#### Study Population

Participants’ consent was obtained prior to genotyping or cognitive testing and they were told they could withdraw from the study at any time. Participants were screened for demographics and clinical characteristics (Table 2) and genotyped for their *APOE* status. Only *ε3*/*ε3* and *ε3*/*ε4* carriers were retained for the current study. The *APOE ε4* carrier (*n*=22) and non-carrier (*n*=22) groups were matched for age, gender, education and overall cognition (ACE-III; Table 2). Participants were tested on the SHQ App. The same SHQ measures of spatial navigation were obtained from the population database and the lab cohort.

### 2.3 Genetic Data

#### Saliva collection and DNA extraction

DNA was collected using a Darcon tip buccal swab (Fisher Scientific, Leicestershire, United Kingdom, LE11 5RG). Buccal swabs were refrigerated at 2-4°C until DNA was extracted from the cheek swabs using the QAIGEN QIAamp DNA Mini Kit (QAIGEN, Manchester, United Kingdom, M15 6SH). DNA was quantified by analysing 2 μL aliquots of each extraction on a QUBIT 3.0 Fluorometer (Fisher Scientific, Leicestershire, United Kingdom, LE11 5RG). Successful DNA extractions were confirmed by the presence of a DNA concentration of 1.5μg or higher per 100μg AE buffer as indicated on the QUBIT reading.

#### Real-time PCR APOE SNP genotyping assay

PCR amplification and plate read analysis was performed using Applied Biosystems 7500 Fast Real-Time PCR System (Thermo Fisher Scientific, Ashford, United Kingdom TN23 4FD). TaqMan Genotyping Master Mix was mixed with RT-PCR SNP Genotyping Assays to determine the 112 T/C (rs429358) *APOE ε4* polymorphism and 158 C/T (rs7412) *APOE ε2* polymorphism.

### 2.4 Statistical analysis

The data were analysed using SPSS (Version 23) and R (Version 1.0.153). All navigation performance data were transformed to log-scale using natural logarithm (log_e_) due to lack of normality in original scale. Next, linear mixed effect models were used to test the effects of sex, age and APOE status on the log-transformed navigation performance scores. Linear mixed effect models with subject-level random effects were used to take account of possible correlation between repeated measures from multiple levels of the game for the same player.

The outcome variables for allocentric wayfinding performance include the ‘‘distance travelled*’’* score and the ‘‘duration’’ score, which reflect the distance covered and the time taken to complete levels during wayfinding levels respectively. Participants completed three wayfinding levels that increased in difficulty sequentially (specifically SHQ levels 6, 8, and 11). ‘‘Flare accuracy*’’* was calculated as number of stars gained and was utilised as an outcome variable for path integration performance (specifically SHQ level 9 and 14).

In the benchmark data, the independent effects of sex and age on outcome variables distance, duration and flare accuracy were the main variables of interest, with level of the game as a control variable. Similarly, for the lab cohort fixed effects included sex, age, level of the game and *APOE* status. To validate the mixed effects model and explore which levels are driving the results, fixed effects linear regression models with age, sex and *APOE* status (for the lab-based cohort) were fitted for each of the SHQ levels analysed. Student’s t-tests, analyses of variance (ANOVAs) and chi-square tests to compare groups were also used where appropriate. Sex and APOE interaction effects were included in the analysis.

## 3. Results

### 3.1 Descriptive Statistics

The age and sex distribution for the benchmark population and for the lab-based cohort are presented in Table 1. Both populations were within the 50-75 age range. Further, the *APOE ε4* carriers did not differ from non-carriers on medical status or general cognitive abilities (Table 2). Importantly, both groups were statistically similar on their episodic memory performance on the ACE-III.

**Table 1.**
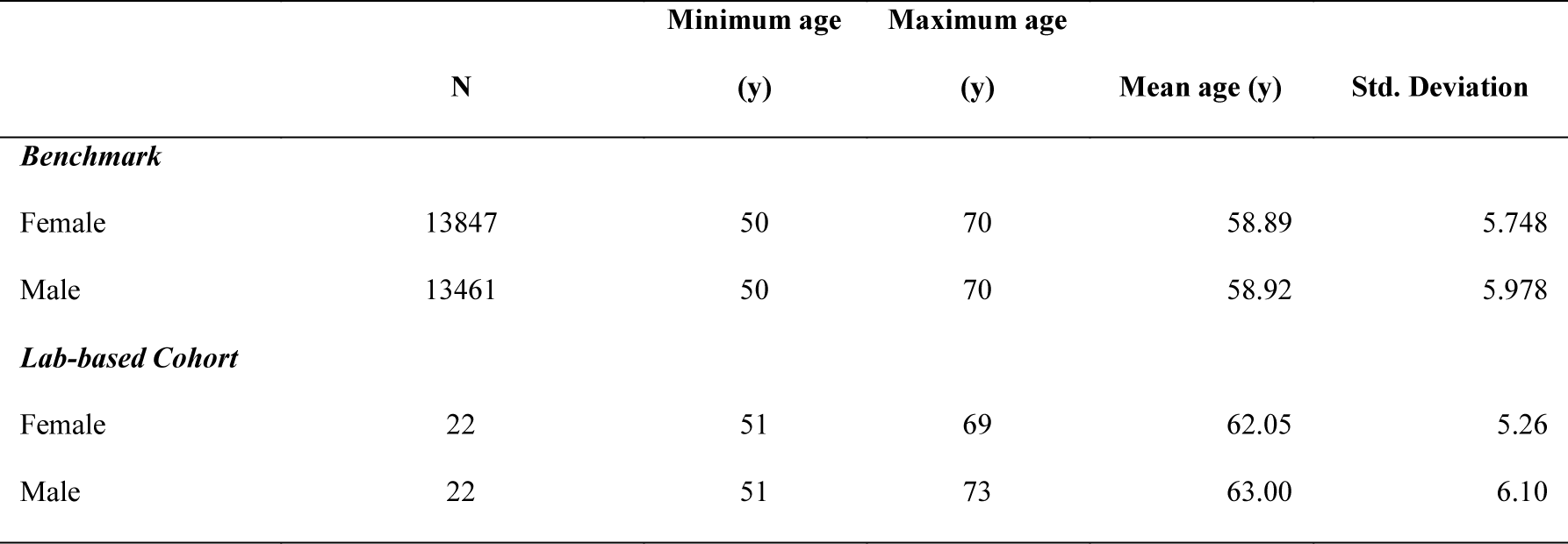
Sex and gender for population and lab-based cohort

**Table 2.** Demographic and clinical outcome measures of the lab-based cohort

| Measure |  | e3/e3 | e3/e4 |  |  |
| --- | --- | --- | --- | --- | --- |
| *General* |  |  |  |  |  |
| *Characteristics* |  |  |  | **Phi/Cramer's V** | **p value** |
| **Sex** | Male | 11 | 11 | 0.099 | p > 0.5 |
|  | Female | 11 | 11 |  |  |
| **Marital Status** | Single | 6 | 4 | 0.189 | p > 0.5 |
|  | Married | 16 | 18 |  |  |
| **Education** | Average Years | 16 | 15 | 0.103 | p > 0.5 |
| **Blood pressure** | Normal (missing 6 subjects) | 18 | 12 | 0.336 | p > 0.5 |
|  | Hypertension | 4 | 4 |  |  |
| **Total cholesterol** | Normal (missing 6 subjects) | 18 | 13 | 0.33 | p > 0.5 |
|  | Medically managed | 4 | 3 |  |  |
| **Family History of AD** | None | 12 | 11 | 0.133 | p > 0.5 |
|  | 1 parent | 6 | 7 |  |  |
|  | 2 parents / 1 early onset | 4 | 4 |  |  |
| *Neuropsychology* |  |  |  | **t** | **p value** |
| **ACE-III \*** | Total score | 93.267 (4.39) | 92.57 (3.6903) | 0.151 | p > 0.5 |
| **Attention** |  | 17.1 (1.1) | 16.6 (1.9) | 1.002 | p > 0.5 |
| **Memory** |  | 24.0 (2.82) | 24.0 (1.37) | 0.792 | p > 0.5 |
| **Fluency** |  | 12.0 (1.49) | 12.1 (1.62) | 0.246 | p > 0.5 |
| **Language** |  | 25.5 (0.87) | 25.5 (0.71) | 0.373 | p > 0.5 |
| **Visuospatial** |  | 14.6 (1.09) | 14.6 (1.17) | -0.074 | p > 0.5 |

Blood pressure and cholesterol were considered abnormal if participant was receiving medication to lower levels. If no medical assistance
was required levels were considered normal. Education includes years at primary, secondary, diploma, undergraduate and or PhD level.
Highly responsible or intellectual occupation was defined as CEO of a large company, judge, university professor, top manager, politician, surgeon, etc. Standard deviation in parentheses. ACE-III \* (Addenbrooke's Cognitive Examination-III)

### 3.2 The impact of age and sex on wayfinding for the benchmark population

In the benchmark database, a linear mixed model revealed significant effects of both sex (regression coefficient [b]=0.13, *p*<0.001, 95% CI= 0.119–0.131) and age (b=0.008, *p*<0.001, 95% CI=0.007–0.008) on wayfinding distance. The coefficient for sex indicates that, independent of age and level of the game, women on average scored 0.13 (in log-scale) higher than men, with higher scores indicating poorer performance when a less efficient route to the checkpoint goal was taken which is reflected in a greater distance travelled within wayfinding levels.

To confirm these gender and age effects, we analysed each wayfinding level separately in a fixed effects linear regression model, resulting in estimated regression coefficients (b) of 0.09, 0.12 and 0.161 for levels 6, 8 and 11, respectively. These coefficients revealed that women travelled 0.09, 0.12 and 0.161 further (in log scale) than men on wayfinding levels. Regression coefficients for age revealed distance scores increase by 0.03, 0.04, and 0.05 (in log scale) on levels 6,8,11 respectively for every 5-year in increase in age. Each of the regression coefficients were significant at *p*<0.001

Similarly, effects of both sex (b=0.14, *p*<0.001, 95% CI= 0.134–0.152) and age (b=0.01, *p*<0.001, 95% CI=0.01–0.012) on wayfinding duration score were found to be statistically significant in a mixed effects model. When fixed effects linear regression models were run on individual levels, regression coefficients of 0.103, 0.128, and 0.198 for sex effects and 0.009, 0.011, and 0.012 for age effects on levels 6, 8 and 11 were found (*p<*0.001). Women’s scores were 0.103, 0.128, and 0.198 points (in log scale) higher on average than men’s scores on levels 6, 8, and 11 respectively, with higher scores indicating longer time taken to complete the level. Coefficients 005, 0.05, and 0.06 for age on levels 6, 8 and 11 respectively, show a steady increase in duration taken to complete the levels for every 5 additional years

### 3.3. The impact of age and sex on path integration (flare accuracy) for the benchmark population

In the population database, a linear mixed model revealed significant effects of both sex (b=-0.09, *p*<0.001, 95% CI=-0.10–-0.082) and age (b= −0.003, *p*<0.001, 95% CI=-0.000–-0.004) on flare accuracy. Thus, women score 0.09 points (on log scale) lower on flare accuracy than men, with lower scores indicating poorer performance. Furthermore, being 5 years older lead to a 0.003 points reduction in average score. For levels 9 and 14, linear regression models estimated individual regression coefficients of −0.071 and −0.116 respectively for sex, showing that differences across sexes increased with level difficulty (consistent with wayfinding levels above) and that women consistently scored lower than men across both flare levels. Regression coefficients of −0.001 (*p=*0.013) and −0.006 for age (*p*<0.001) reveal that performance declines with age on flare accuracy, which is consistent with performance on wayfinding (Figure 1. ii).

### 3.4 The impact of age, sex and APOE on wayfinding in the lab-based cohort

In the lab-based cohort, an effect of *APOE* on wayfinding distance (b=0.13, *p*=.02, 95% CI=0.024–0.239) but no effects of sex (b=0.05, *p*=0.36, 95% CI=-0.057–0.159) or age (b=- 0.001, *p*=0.70, 95% CI=-0.012–0.008) were found in a mixed effect linear model. Specifically, *ε3*/*ε4* carriers scored 0.13 points (on log scale) higher than *ε3*/*ε3* carriers on wayfinding distance, with higher scores indicating poorer performance. The effect of *APOE* was driven by *level 11* (*APOE* effects of b=0.16, *p=*0.027 in a linear regression with sex and age as other predictors), as there were no significant effects of the gene, age or sex on the easier levels 6 and 8 (*p=*>0.05) (Figure 2).

**Figure 2.**
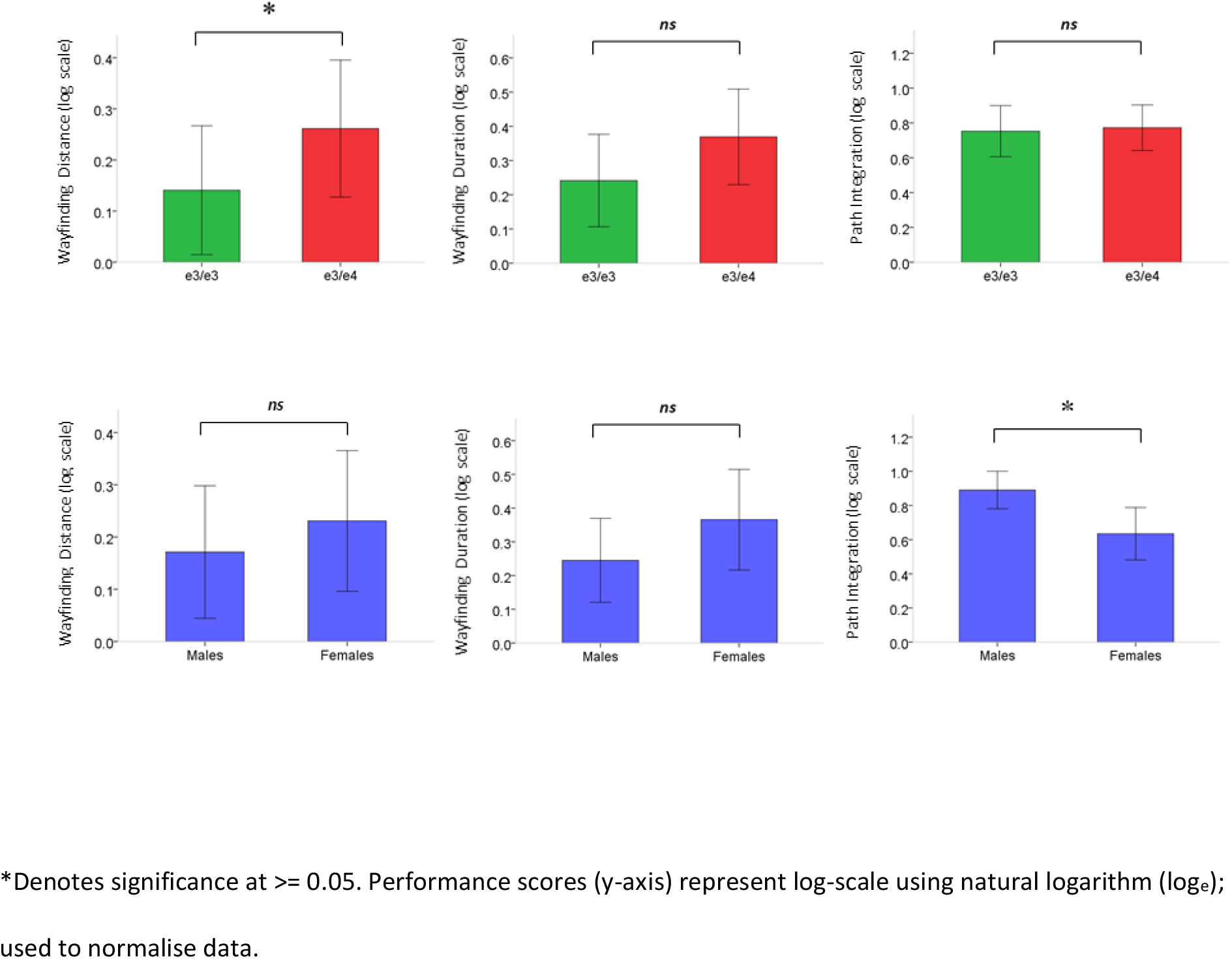
*APOE* genotype and sex effects on wayfinding distance, wayfinding duration and path integration performance for the lab-based cohort.

No significant effects of either *APOE* (b=0.12, *p*=0.94, 95% CI=-0.0172–0.257), sex (b=0.13, *p*=0.66, 95% CI=-0.005–0.269) or age (b=-0.001, *p*=0.78, 95% CI=-0.014–0.011) were found on the wayfinding duration scores. Although *APOE* and sex effects did not reach significance, coefficients for the contribution of sex (b=0.13) and *APOE* (b=0.12) were of similar magnitude to the coefficients for the sex effects in the population data analyses (see Figure 2). In fact, a linear regression model revealed significant sex effects (b=0.22, *p*=0.03) on the harder *level 11*. Finally, no interaction effects of sex and *APOE* on distance travelled or duration taken during wayfinding levels were found.

### 3.5 The impact of age, sex and APOE on path integration (flare accuracy) in the lab-based cohort

A significant effect of sex on flare accuracy scores was found in the lab cohort (b=-0.28, *p*=0.009, 95% CI=-0.473–-0.08) using the mixed effects approach, with women scoring −0.28 points lower than men on flare accuracy. This result was found to be driven by level 14 (b= - 0.44, *p=*0.005) rather than level 9 (b= −0.21, *p=*0.12). There was no effect of the *APOE* genotype (b=0.03, *p*=0.703, 95% CI=-0.159–0.236) or age (b=-0.001, *p*=0.972, 95% CI = −0.018–0.017) on flare accuracy (Figure 2). See Table 3 for all regression coefficients mentioned above.

**Table 3.** Regression coefficients for mixed effects linear models.

| Independent Variables | Predictors | b coefficient | Std. Error | P value |
| --- | --- | --- | --- | --- |
| *Population Model* |  |  |  |  |
| Wayfinding Distance | Gender | **0.13** | 0.003 | **p<0.001** |
|  | Age* | **0.008** | 0.0003 | **p<0.001** |
| Wayfinding Duration | Gender* | **0.14** | 0.004 | **p<0.001** |
|  | Age* | **0.011** | 0.0004 | **p<0.001** |
| Path Integration | Gender* | **-0.09** | 0.004 | **p<0.001** |
|  | Age* | **-0.003** | 0.0004 | **p<0.001** |
| *Lab-based Cohort Model* |  |  |  |  |
| Wayfinding Distance* | Gender | 0.05 | 0.055 | 0.36 |
|  | Age | -0.002 | 0.005 | 0.70 |
|  | **APOE\*** | **0.13** | **0.055** | **0.02** |
| Wayfinding Duration | Gender | **0.13** | 0.067 | **0.07** |
|  | Age | -0.002 | 0.006 | 0.78 |
|  | APOE | **0.12** | 0.070 | **0.09** |
| Path Integration | **Gender\*** | **-0.28** | 0.101 | **0.01** |
|  | Age | -0.0003 | 0.009 | 0.97 |
|  | APOE | 0.04 | 0.101 | 0.70 |

Fitted parameters from mixed effects linear models. Bold indicates significant or approaching significant effects.

### 3.6 Direct comparison of benchmark population and lab-based cohorts

The lab cohort revealed that *APOE ε4* carriers performed worse on *level 11* of SHQ than *ε4* non-carriers. To investigate the robustness of this finding, individuals with known genotype were compared to the benchmark population cohort. The rationale behind this post-hoc analysis was that, based on existing epidemiological studies of *APOE* genotype prevalence, the majority of the population cohort (~75%) are non-*APOE ε4* allele carriers and hence should be in its performance more similar to the *ε3* carriers than the *ε4* carriers in the lab cohort.

Despite sample size inequality between the population (n=27,308) and lab (n=44) cohorts, a student’s t-test assuming unequal variances uncovered significant differences between *ε4* carriers and the benchmark population mean (t_21.1_=-2.373, *p=*0.027). At the same time, no significant differences between *ε3* carriers in the subpopulation and the primarily *ε3* allele carrying benchmark population mean were found (t_21.1_=0.471, *p=*0.643) (Figure 3.i). No differences were observed in *level 11* duration scores between the *ε3*, *ε4* carriers and the benchmark population (*p*=0.23) although women spent significantly longer on solving *level 11* than men from the lab cohort (Figure 3.ii).

**Figure 3.**
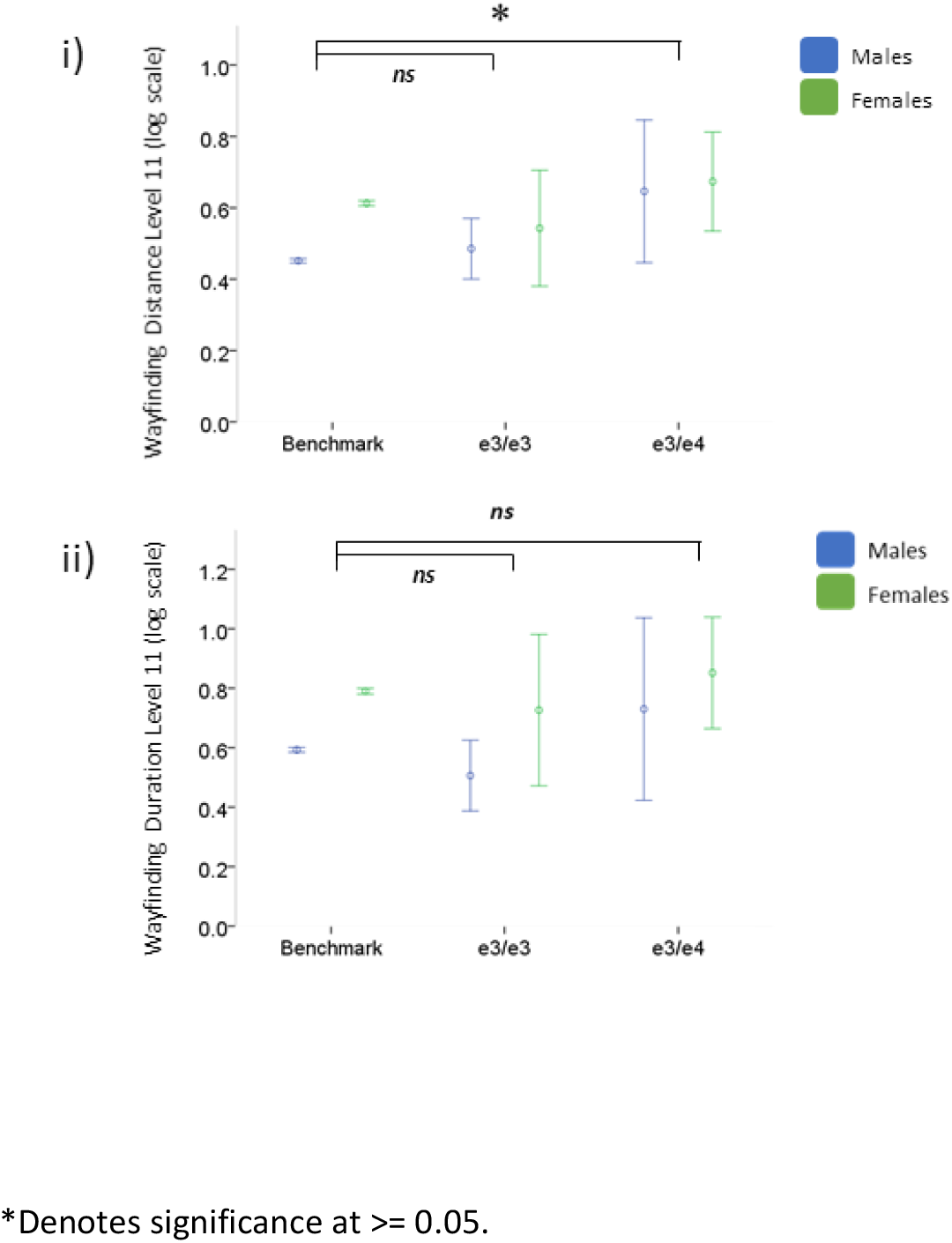
Direct comparison of benchmark population and lab-based cohorts (e3/e3 and e3/e4 groups) on level 11 i) wayfinding distance and ii) wayfinding duration.

## 4. Discussion

Our results clearly show that both sex and *APOE* status impact on spatial navigation performance in midlife ageing cohorts, while age had a surprisingly much weaker effect on performance. More specifically, the population data showed a consistent significant effect of sex on spatial navigation performance for both wayfinding and path integration.

Interestingly, men outperformed women on both spatial process. Age explained less variance in spatial performance than sex, primarily predicting wayfinding ability. Likely due to reduced statistical power due to a small sample size in the lab cohort, we only observed sex effects for path integration and not for wayfinding. More importantly, adults who have a higher genetic risk for AD due to *APOE ε4* presence showed a specific, sex-independent deficit in spatial navigation for wayfinding. This deficit was confirmed by comparing wayfinding performance of *ε3* and *ε4* carriers from the lab cohort against the population cohort primarily consisting of *ε3* carriers.

The effects of sex and age on spatial navigation performance within the benchmark population are consistent with previous experiments [15–21]. The influence of age may be linked to age-related cerebral changes within the dentate gyrus and CA3 region of the hippocampus which would naturally result in age having a greater effect on allocentric wayfinding ability than egocentric path integration ability[2]. However, explanations for the influence of sex on spatial navigation skill are multifaceted and both social and biological theories have been proposed. It is interesting that within the current study, the pattern of age-related decline on wayfinding and path integration did not appear to differ between men and women, although some previous studies suggest a more pronounced decay in spatial memory is present in women [22,23]. The influence of sex was greater than age in both cohorts, supporting that assumption that the negative impact of age on spatial navigation emerges after the age of 70 [9]. Further, the predictive value of age on wayfinding was almost triple the predictive value of age on path integration in the population data, consistent with evidence that age-related deficits appear first in wayfinding-based allocentric processing and only later in path integration-based egocentric processing [7,24,25]

As mentioned above, in the lab cohort sex effects were observed for path integration but not for wayfinding. Instead, *APOE* status was the greatest predictor of wayfinding performance, showing that genetically at-risk adults travelled a longer distance particularly in the harder wayfinding levels. This finding nicely dovetails results by Kunz and colleagues (2015) who showed that *APOE* leads to a reduced preference to navigate in the centre of an environment. Kunz et al attributed these genetic differences to reduced grid-cells representations in the entorhinal cortex of *ε4* carriers. Although fMRI data is needed to elucidate the neural mechanisms behind the altered spatial trajectory present in our genetically at-risk group, our behavioural data suggests that at-risk healthy participants show such subtle trajectory changes in SHQ as they covered a greater distance and therefore took a less efficient route to the check-point goal during allocentric wayfinding. This suggestion clearly needs further substantiated. Although the impact of *APOE ε4* on spatial navigation did not appear to be greater in women, women’s overall spatial ability was poorer than men’s, with stable sex differences on path integration found across both databases. This mirrors previous research showing that wayfinding does differ among males and females and therefore investigating the interaction between *APOE* and sex in a larger cohort is required.

On a clinical level, sex differences found on the population level demonstrates that diagnostically sex needs to be accounted for when considering spatial navigation as a cognitive marker in pre-symptomatic individuals. Specifically, cut-off scores to determine whether deficits are present or not should be based of different normative data for women and men. Although there are potentially unknown reasons why a person may perform below average, if sex differences are controlled for and a person still performs poorly, this should raise a question regarding the persons *APOE* status, which can then elucidate their future risk of developing AD pathophysiology. Based on this information and stratification, prevention and treatment plans can be tailored better to those individuals at high risk of developing AD in the population.

Despite these exciting findings, our study has several shortcomings. Online data collection has often the reputation of suffering from selection bias (i.e. people engaging in online tasks are more likely to self-select). Our SHQ results are unlikely to be confounded by selection bias as the sample size exceeded 27,000 people and thus only a small percentage of participants would fall likely under this category. Further, our validation of the population data scores against the lab cohort of *ε3* carrier scores indicates that the population sample is representative of a non-risk cohort. In terms of the lab cohort, ideally, we would have had a similar sample size to the population cohort to corroborate our genetic findings. But this is clearly not practical as the smaller sample size might have led to lower statistical power and hence no significant findings for sex in the wayfinding conditions in the lab cohort. It would be therefore important to replicate these findings in the future in larger genotyped cohorts. Still, the sample sizes for the *APOE* cohort were of similar magnitude to other high impact navigation studies in the field.

Taken together, these findings suggest that sex and APOE genotype information can be used to personalise diagnostic and treatment approaches in the future, and further that the utility of digital spatial navigation tests to detect high-risk individuals in the population is promising.

## Acknowledgments

The authors wish to thank Deutsche Telekom for funding the development of Sea Hero Quest. In addition, we would like to thank Deutsche Telekom and Alzheimer’s Research UK for their funding support in the analysis of the Sea Hero Quest data. We thank the Glitchers Limited for the game production of Sea Hero Quest. We also wish to extend a thank you to all participants who devoted their time to this research.

